# Spatial variation in introgression along a toad hybrid zone in France

**DOI:** 10.1101/746073

**Authors:** I. van Riemsdijk, J.W. Arntzen, G. Bucciarelli, E. McCartney-Melstad, M. Rafajlović, P.A. Scott, E. Toffelmier, H. B. Shaffer, B. Wielstra

**Affiliations:** Naturalis Biodiversity Centre, Leiden, the Netherlands; Institute of Biology Leiden, Leiden University, Leiden, the Netherlands; Department of Ecology and Evolutionary Biology, UCLA, Los Angeles, California; La Kretz Center for California Conservation Science, Institute of the Environment and Sustainability, UCLA, Los Angeles, California; Department of Marine Sciences, University of Gothenburg, Gothenburg, Sweden

**Author notes:** Correspondence: Isolde van Riemsdijk.

**Keywords:** Asymmetric introgression, barrier genes, *Bufo bufo*, *Bufo spinosus*, replicate transects, cline coupling

## Abstract

The barrier effect is a restriction of gene flow between diverged populations by barrier genes. Restriction of gene flow and asymmetric introgression over multiple transects indicates species wide (genetic) adaptations, whereas transect-specific barrier loci may indicate local adaptation to gene flow. Asymmetric introgression can be caused by selection, hybrid zone movement, asymmetric reproductive isolation, or a combination of these. We study two widely separated transects (northwest and southeast France) for the 900 km long hybrid zone between *Bufo bufo* and *B. spinosus* toads, using ~1200 markers from restriction-site associated DNA (RAD) sequencing data. Genomic and geographic clines were used to identify outlier markers which show restricted or elevated introgression. Twenty-six barrier markers are shared between transects (the union of 56 and 123 barrier markers identified in each transect), which is more than would be expected by chance. However, the number of barrier markers is twice as high in the southeast transect. In the northwest transect a high amount of (asymmetric) introgression from *B. spinosus* into *B. bufo* is consistent with hybrid zone movement or asymmetric reproductive isolation. In the southeast transect, introgression is symmetric and consistent with a stable hybrid zone. Differences between transects may be related to genetic sub-structure within *B. bufo*. A longer period of secondary contact in southeast France appears to result in a relatively stronger barrier effect than in the northwest. The *Bufo* hybrid zone provides an excellent opportunity to separate a general barrier to gene flow from local reductions in gene flow.

## Introduction

Hybrid zones provide an opportunity to study both the processes involved in, and the outcomes of, speciation (Hewitt, 1988). Barrier genes, defined as genomic regions that restrict gene flow and introgression between hybridizing populations, play a key role in speciation (Ravinet et al., 2017). Barrier genes can comprise genes involved in divergent ecological selection, mate choice, and genetic incompatibilities (Ravinet et al., 2017). Barrier genes can create genomic heterogeneity, with relatively strong differentiation of these genes themselves as well as the genomic regions surrounding them, which may prohibit complete merging of the parental populations (Abbott et al., 2013; Barton, 2013; Ravinet et al., 2017).

The barrier effect is a reduction of effective migration rate of genetic material between populations, relative to the dispersal of the individuals carrying those genes (Ravinet et al., 2017). Barrier genes and their linked loci (together referred to as ‘barrier markers’ here) are expected to show relatively steep transitions at species boundaries (Gompert, Parchman, & Buerkle, 2012; Butlin & Smadja, 2018). This barrier effect is reinforced when the gene frequency clines of multiple barrier genes become geographically coincident in a hybrid zone, a phenomenon referred to as cline coupling (Butlin & Smadja, 2018). When the same markers show such steep and co-distributed transitions along multiple, geographically distant transects across a hybrid zone, it is most likely that the two hybridizing species evolved a barrier effect across their entire species’ range (Teeter et al., 2009; Larson, Andrés, Bogdanowicz, & Harrison, 2013; Harrison & Larson, 2014; Larson, White, Ross, & Harrison, 2014).

Patterns of introgression can be indicative of different types of selection. Asymmetric introgression in hybrid zones, where gene flow is more pronounced in one direction than in the other, can be caused by hybrid zone movement, positive selection, asymmetric reproductive isolation, or a combination of these. Hybrid zone movement occurs when one species outcompetes or out-disperses the other, and can cause elevated introgression of many neutral markers in the wake of the movement (Gay, Crochet, Bell, & Lenormand, 2008; Excoffier, Foll, & Petit, 2009; Wielstra, Burke, Butlin, & Arntzen, 2017; Wielstra, Burke, Butlin, Avcı, et al., 2017; Wielstra, 2019). Asymmetric pre- or postzygotic isolation involves the successful reproduction of only certain combinations of individuals (e.g. hybrids can only backcross with one of the parental species), and most of the introgression takes place on the side of the hybrid zone where backcrossing is the most successful (Haldane, 1922; Hewitt, 1975; Barton, 2001; Devitt, Baird, & Moritz, 2011). As hybrid populations receive more genes from the side of the hybrid zone where reproduction is most successful, asymmetric reproductive isolation can be a competitive advantage, and thus result in hybrid zone movement (Buggs, 2007). Adaptive introgression, on the other hand, would typically concern only one or a few markers (Barton & Hewitt, 1985; Barton, 2001).

The use of multiple transects across the same hybrid zone provides the ability to distinguish between general and local patterns of gene flow, and thus to identify pervading processes causal in reproductive isolation. We study two unique transects at opposite ends of a ca. 900 km long hybrid zone between the common toad, *Bufo bufo* (Linnaeus, 1758) and the spined toad, *B. spinosus* Daudin, 1803, which runs diagonally across France from the Atlantic coast in the north to the Mediterranean Sea in the south (Fig. 1; Arntzen et al., 2018). Given the low dispersal of both species, the *Bufo* hybrid zone provides an opportunity to test the consistency of patterns of restricted gene flow and elevated directional gene flow in independent transects. Previous studies on the individual transects suggested that weak asymmetric introgression may occur in both sections (Arntzen, de Vries, Canestrelli, & Martínez-Solano, 2017; van Riemsdijk, Butlin, Wielstra, & Arntzen, 2018). Expanding on this work, we use ~1200 nuclear markers, two to three orders of magnitude more than used in previous studies, to assess genome wide patterns on introgression in this hybrid zone. Assuming that reproductive isolation is consistent throughout the hybrid zone, we hypothesise that the same barrier genes are consistently reducing gene flow between the two species in both transects.

**Figure 1:**
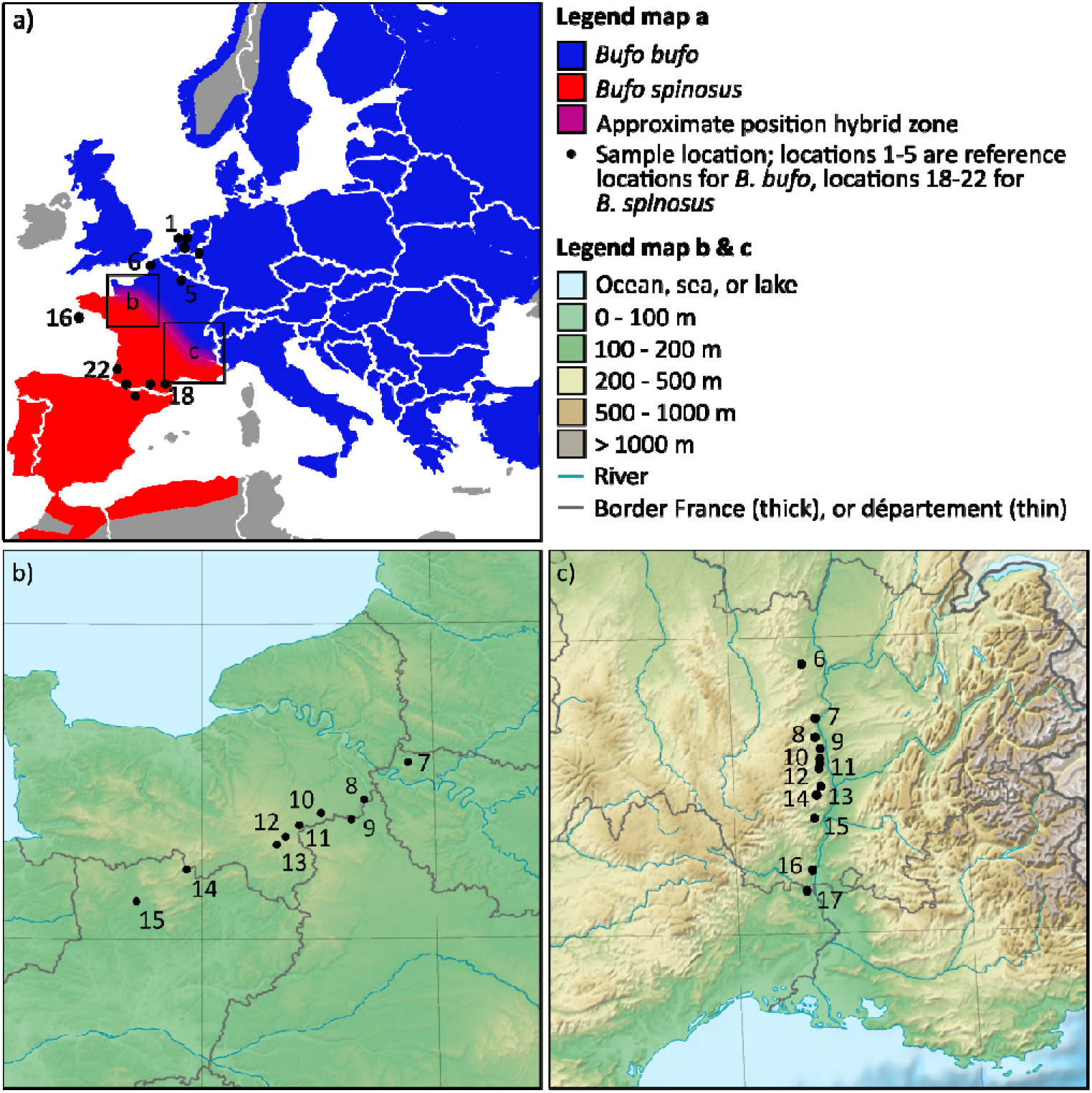
Overview map with a) the distribution of *Bufo bufo* and *B. spinosus*, with small squares indicating the locations of map b) for transect one in northwest France (sample locations 6-16), and map c) for transect two in southeast France (sample locations 6-17). The base map for panel b and c was downloaded from https://www.mapsland.com

## Material and Methods

### 3RAD sequencing

DNA extracts of 387 individual toads were reused from previous research (Arntzen et al., 2016, 2017, 2018, in prep.). This included five reference sample sites each of presumably pure *B. bufo* and *B. spinosus*, 11 sample sites in transect one in northwest France, and 12 sample sites in transect two in southwest France (Table S.1.; Fig. 1). For three presumably pure individuals of each species, libraries were prepared in triplicate (6 individuals x 3 = 18) to assess genotyping error rate after assembly (which was estimated to be 0.5%; Appendix 1). The 3RAD method (Graham et al., 2015; Glenn et al., 2016; Hoffberg et al., 2016; Bayona-Vásquez et al., 2019) was used to obtain reduced representation genomic libraries. Two restriction enzymes (CLA-*I* and Sbf-*I*) were used to cut 50 ng of genomic DNA from each sample, while a third enzyme (MSP-*I*) was added to cleave and eliminate phosphorylated adapter-adapter dimers. Internal barcodes were ligated to the resulting sticky ends and external Illumina iTru5 and iTru7 primers, differing by ≥ 3 bp, were added to the internal barcodes via an indexing PCR reaction (Glenn et al., 2016; Hoffberg et al., 2016; Bayona-Vásquez et al., 2019). For PCR details see supplementary data (High-Throughput 3RAD Protocol) in Bayona-Vásquez et al. (2019).

Upon individual library completion, DNA concentrations of seven libraries contained less than the required concentration for equimolar pooling, even after repeating the library preparation twice, thus the total volume of the library was included in the pool. All other libraries were combined to achieve equimolar concentrations in the final pool. The fragments in the pooled library were size selected for a range of 340-440 bp using a Pippin Prep (Sage Science Inc. Beverly, MA), quantified using intercalating dye on a Victor multilabel plate reader, and were sequenced on two lanes of an Illumina HiSeq 4000 PE100 at the Vincent J. Coates Genomics Sequencing Laboratory in Berkeley, CA, USA.

### Data clean-up and assembly

The average read count per sample was ~1.5 million paired-end reads (maximum 4.2 million reads, Table S.2). Twelve samples had low raw read quantities (near or below 0.5 million reads), including the seven samples with low DNA concentration, and were excluded. Cutadapt v.1.14 (Martin, 2011) was used in three steps to remove 5’ and 3’ primers for each internal barcode combination, remove the Illumina standard adapter, and carry out a read quality control. Subsequently, ipyrad v.0.7.3 (Eaton, 2014) was used to assemble the reads. Settings were: minimum read depth of six, maximum of eight heterozygous bases allowed per consensus sequence, and heterozygous sites allowed across a maximum of 50% of the samples (details in the supplements).

### Population structure

Using a single randomly selected SNP per RAD fragment from the 50% dataset (which has 47.5% missing data) and the 90% dataset (which has 2.3% missing data), we first ran a principal component analysis (PCA) to visualize genomic variation across both transects. We used the package ‘adegenet’ v.2.1.1 in R (Jombart, 2008; Jombart & Ahmed, 2011), and based the PCA on allele frequencies, replacing missing data with the mean of the total dataset.

We further quantified population structure with Structure v.2.3.4 (Pritchard, Stephens, & Donnelly, 2000) with the same datasets, for each transect separately. For both data sets, ten independent Structure runs for two to ten genetic clusters (*K*) were completed with a burn in of 10,000 MCMC steps followed by 25,000 MCMC steps under the admixture model using StrAuto (Chhatre & Emerson, 2017). Convergence of the results was checked by investigating log likelihood and admixture proportion stability (Benestan et al., 2016). The results were summarized with CLUMPAK (Kopelman, Mayzel, Jakobsson, Rosenberg, & Mayrose, 2015). The optimal value of *K* was determined with the Evanno method (Evanno, Regnaut, & Goudet, 2005). Structure results were visualised using the R package POPHELPER (Francis, 2017).

### Diagnostic SNP selection

Diagnostic SNPs were determined based on the genotypes of the 37 *B. bufo* and 20 *B. spinosus* individuals from the reference sample sites (Fig. 1; Table S.1, custom R script). In order to verify whether there was bias from taxon-specific patterns of missing data, missing data matrices were constructed with the 50% and 90% data sets, showing that data missingness is slightly biased towards *B. spinosus*, which is expected, as it is the more genetically diverse species of both (Fig. S.1) - so comparatively higher levels of allelic drop-out from mutated restriction sites would be expected in this species. To avoid these possible null-alleles, we removed all SNPs for which a locus was missing in all samples from the reference sample sites of a species from the dataset with a maximum of 50% missing SNPs per individual. We then selected all bi-allelic SNPs that were fixed for alternative homozygous variants in the set of reference samples of each species. Finally, we selected one random diagnostic SNP per fragment, resulting in 1,189 diagnostic SNPs. The percentage of missing data in the dataset after diagnostic SNP selection was 25.7%.

### Hardy-Weinberg equilibrium

We tested for signals of non-random mating success for each sample site in the dataset with diagnostic SNPs by calculating heterozygote excess and deficit from Hardy-Weinberg equilibrium with the R package ‘genepop’ based on the program GENEPOP v.1.0.5 (Rousset, 2008). Instead of the conservative Bonferroni correction, which accounts for the number of tests performed in total, independence of tests was accounted for within markers (Pc for N=1,189; Rice 1989; Narum 2006). Many markers with a significance level uncorrected for repeated testing (P < 0.05) in the hybrid zone populations for both heterozygote excess and deficit were present, but only 14 markers had a significant heterozygote deficit after correction (Table S.3). These markers were excluded in the HZAR geographic cline fitting analysis and admixture linkage disequilibrium calculations (see below).

### Bayesian genomic cline outlier detection

To study genome-wide variation of introgression among admixed individuals we used the Bayesian genomic cline model as implemented in the software BGC (Gompert & Buerkle, 2011, 2012; Gompert et al., 2012). The Bayesian genomic cline model is based on the probability that an individual with a certain hybrid index (HI) inherited a gene variant at a given locus from one species (φ; in this case *B. bufo*) or the other (1 − φ; *B. spinosus*). The probability of *B. bufo* ancestry relative to expected (represented by the HI) is described by cline parameter α. A positive α indicates an increase in the *B. bufo* ancestry probability and a negative α indicates a decrease. The cline parameter β measures the genomic cline rate based on ancestry for each locus. A positive β indicates an increased transition rate from a low to high probability of *B. bufo* ancestry as a function of the HI, which implies there are less heterozygotes for the marker than expected based on the HI, whereas a negative β indicates a decrease in the transition rate, which implies there are more heterozygotes than expected based on the HI (Gompert & Buerkle, 2011; Parchman et al., 2013). When more markers are an outlier for α in one direction than the other, this points to genome wide asymmetric introgression. When a marker is a negative outlier for cline parameter β, it is a candidate for a barrier marker, especially when the geographic cline is narrow and located in the centre of the hybrid zone.

The input files for parental genotypes included only individuals from the reference sample sites, and the input file for admixed genotypes included individuals with an average admixture proportion (Structure Q score) between 0.05 and 0.95, treated as a single population (Fig. S.2). A single MCMC chain was run for 75,000 steps and samples were taken from the posterior distribution every 5^th^ step, following a burn-in of 25,000 steps. Convergence was assessed (Fig. S.3), and we tested for outlier loci using ‘estpost’ to summarise parameter posterior distributions (Gompert & Buerkle, 2011). Outlier loci were established based on 99.9% confidence intervals of parameters.

### Geographic cline analysis

Classic geographic equilibrium cline models were fitted using the R package ‘HZAR’ (Derryberry, Derryberry, Maley, & Brumfield, 2014) for all diagnostic SNPs. For transect one, sample sites 6 and 16, which are distant from the main axis of the transect, were removed from the dataset. To determine distance between the sample sites, a custom R script was used, and the directions of the transect axes were the same as in previous publications (Arntzen et al., 2016, 2017). Transect two is situated next to the Rhone river, which is considered a barrier to dispersal, and therefore the only logical direction of the transect is parallel to the river. The shapes and positions of many clines can be summarised in the expected cline, which can be represented by the HI (Polechová & Barton, 2011; Fitzpatrick, 2012). We thus also fitted clines for the HI of all non-outlier markers as determined by BGC, as well as the HI of all heterozygote deficiency outliers (β > 0), to be able to compare the general shape and positions of these marker categories in a geographical setting. For example, the heterozygote deficiency outliers are expected to be geographically steep clines.

Thirty maximum likelihood estimation searches were performed with random starting parameters, followed by a trace analysis of 60,000 generations on all models with a delta Akaike information criterion corrected for small-sample-size (dAICc) < 10. Fifteen model variants were based on all possible combinations of trait intervals (allele frequency at the ends of the transects; three types) and tail shape (five types). Even though markers were restricted to be diagnostic, cline shapes sometimes were better described by clines with allele frequencies at the end of the cline different from zero or one. Convergence was visually assessed in trace plots (see supplemental material). To provide a measure of cline symmetry, we used a custom R script to estimate the area underneath the cline tail towards the left (Q_left_) and the right (Q_right_) of the hybrid zone centre, up to the point where the HI reached a value of 0.05 or 0.95 (Fig. S.4, supplements).

### Admixture linkage disequilibrium and effective selection

To assess the most recent inflow of parental genotypes, linkage disequilibrium can be used. The first generation offspring of two diverged species will be heterozygous for all fixed differences, resulting in complete admixture linkage disequilibrium (D’), rather than the usual linkage disequilibrium resulting from selection, assortment, mutation, or drift (Barton & Gale, 1993; Baird, 2015). Recombination during reproduction breaks down D’, whereas migration of parental (pure) individuals increases D’ (Barton & Gale, 1993). When gene flow into the hybrid zone is symmetric, the peak of D’ in the hybrid zone centre follows a Gaussian curve (Gay et al., 2008). Under hybrid zone movement, the peak is predicted to shift ahead of the movement to the side of the hybrid zone, opposite to the tail of neutral introgression where recombination has broken down the peak already (Gay et al., 2008; Wang et al., 2011). The peak is expected to be more coincident with the tail of introgression in a case of asymmetric reproductive isolation (Devitt et al., 2011). The position of the peak of D’ thus may be used to support the underlying process of asymmetric introgression.

Average effective selection on a locus (s*) is the selection pressure on a locus at the zone centre due to direct selection and association with other loci. Admixture linkage disequilibrium (*D*’) based on the variance in hybrid index, which in turn allows the calculation of lifetime dispersal distance weighted for pre- and post-metamorphosis (σ), and s* following Barton & Gale (1993), were calculated using scripts from van Riemsdijk et al. (2019). The data contained only markers in Hardy-Weinberg equilibrium and markers not indicated as outliers in BGC (832 and 652 loci) for each transect, because these markers represent the presumably neutral portion of the genome (Table 1). We repeated the analysis using only the markers that were heterozygote deficiency outliers (β > 0; 56 and 121 loci) to represent the portion of the genome that experiences the highest barrier effect. Fixed parameters were: a recombination rate of 0.4997, calculated following formula (6) from Macholán et al. (2007), using the number of chiasmata per bivalent for *B. bufo* (1.95; Wickbom, 1945) and the number of chromosomes for *B. bufo* (N = 22), a generation time (sexual maturity) of 2.5 years for *Bufo* at the latitude of the hybrid zone (mean of 3 years in females and 2 years in males; Hemelaar, 1988) and initial secondary contact 8,000 years ago following Arntzen et al. (2016). The width of the hybrid zone was derived from a general sigmoid cline model following HZAR (Derryberry et al., 2014), fitted to the HI, as tail shape is not taken into consideration in these calculations (Barton & Gale 1993). Mean and 95% confidence interval (CI) were based on 1,000 bootstrap replicates of the original genotype dataset (with replacement, maintaining original sample size within sites). Following Gay et al. (2008), we fitted a Gaussian curve through the estimates of *D’* and 95% confidence intervals (CI) for the calculated parameters were derived from the bootstrap data.

**Table 1:** Bayesian genomic cline (BGC) results comparing significant outliers for transect one (T1) and transect two (T2), and the markers which were outliers in both transects (overlap), where significance of outliers is based on the exclusion of 0 in the 99.9% confidence interval (CI). The total number of markers analysed was 1,189.

| Outlier | Biological interpretation | T1 | T2 | Overlap |
| --- | --- | --- | --- | --- |
| $\alpha < 0$ | Directional introgression from *B. bufo* into *B. spinosus* | 151<sup>†</sup> | 185<sup>‡</sup> | 92<sup>§</sup> |
| $\alpha > 0$ | Directional introgression from *B. spinosus* into *B. bufo* | 110<sup>†</sup> | 174<sup>‡</sup> | 49<sup>§</sup> |
| $\beta < 0$ | Heterozygote excess | 50 | 105 | 11<sup>¶</sup> |
| $\beta > 0$ | Heterozygote deficiency | 56 | 123 | 26<sup>§</sup> |

The significance tests are; (1) a Chi-squared comparing the values $\alpha$ outliers of only T1 and T2 to test if there is a significant difference in the number of outliers between the transects. A 2x2 contingency Chi-squared (2) was conducted to test if the overlap between the transects of both $\alpha$ and $\beta$ outliers could be a coincidence, or is unlikely to have occurred under a model of random resampling.
<sup>†</sup>Significant Chi-squared with 6.4406, df = 1, P = 0.0112
<sup>‡</sup>Not significant Chi-squared with 0.3371, df = 1, P = 0.5615
<sup>§</sup>Significant 2x2 contingency Chi-squared with 285.05, 91.629, 81.288, df = 1, P < 2.2e<sup>-16</sup>
<sup>¶</sup>Significant 2x2 contingency Chi-squared with 10.331, df = 1, P = 0.001308

## Results

### Data clean-up and assembly

We assembled two data sets in which the minimum number of individuals that must have data for each locus to be retained was 194 (50% of the samples), and 349 (90%) of the 387 total samples to generate matrixes with different levels of missing data for downstream analyses. These two datasets eventually contained 4,869 loci (39,750 SNPs), and 986 loci (10,535 SNPs), respectively. The PCA plots for the 50% and 90% datasets are highly similar (Fig. S.5). The structure results for both datasets showed that using a dataset with loci for which at least 90% of individuals have data, returned qualitatively similar results as using the dataset with more missingness, but more data (mean delta Structure Q score difference/individual = 0.014; standard deviation = 0.016).

Samples that had libraries prepared and sequenced in triplicate were initially analysed as part of the 50% missingness dataset. This showed that on average, pairwise *de novo* assemblies for each individual contained 79.8 % of the same RAD-loci, and for overlapping loci, differed at only 0.5% of the dataset-wide SNP calls. When manually inspected, the differences of SNP calls were often missed heterozygous calls (e.g. A vs R). Overall, this gives us confidence that our *de novo* assembly parameters strike a balance between read depth and missingness. For example, one would expect, setting higher read depth thresholds would result in fewer missed or erroneous SNP calls, but could also result in higher stochastic missingness of RAD-loci for a given sequencing effort.

### Population structure

The first axis (PC1, 26.8%) of the PCA appears to reflect the genetic difference between pure *B. bufo* in the north (right) and pure *B. spinosus* in the south (left), with hybrids in the middle (Fig. S.5). The second axis (PC2, 2.8%) separates the two transects. In Structure, the preferred number of genetic clusters for transect one was K=3, and for transect two K=2 (Fig. S.2). In both transects, the plot for two genetic clusters reflects differentiation between the two species, with hybrids smoothly transitioning between the two, while the three-cluster model in transect one places hybrids in a group of their own. At both K=2 and K=3, both transects have superficially similar structure, and hybrid individuals never approach being “pure” hybrid at K=3. In other words, hybrids are always shown to have ancestry that is admixed between a ‘hybrid’ population and both parental species.

### Bayesian genomic cline outlier detection

The dataset in the BGC analysis shows little bias in the proportion of observed hybrid indices (HI; Fig. S.6). Transect one has significantly more markers with a reduced probability of *B. bufo* ancestry (α < 0, n = 151) than markers with an increased probability (α > 0, n = 110, χ^2^ test *P* = 0.0112; Table 1), relative to the hybrid index. Transect two has a nearly equal number of markers with an increased or decreased probability of *B. bufo* ancestry (α > 0, n = 174; α < 0, n = 185, χ^2^ test *P* = 0.5615). The number of markers with an outlier β in transect one (heterozygote deficiency β > 0, n = 56 and heterozygote excess β < 0, n = 42) is about half the number detected in transect two (β > 0, n = 123, and β < 0, n = 105). Of the RAD markers which are positive or negative outliers for α or β, 22-61% are also outliers in transect two (last column Table 1). Such overlap is unlikely if the two transects were completely evolutionarily independent (Table 1). For example, the chance of the same 26 markers to act as barrier markers in both transects by chance is close to zero (last row Table 1).

### Geographic cline analysis

We verified that outlier markers for α or β in the BGC analysis are correlated to outlier behaviour in the shape and position of their geographic cline, by plotting significant outliers for the parameters α and β from BGC to the geographic cline parameters centre and width determined with HZAR (Fig S.6). When a marker is an outlier for α, it would be expected that the cline is shifted (centre) or has a different shape (e.g. width), or both, compared to the genomic average (e.g. represented by the HI cline). When a marker is an outlier for heterozygote deficiency (β > 0) it should also show a steep geographic cline, coincident with the HI cline (green clines, Fig. 2, Fig. S.7). We refer to such heterozygote deficiency (β > 0) markers as ‘barrier markers’. Markers that are not genomic outliers (not an outlier for α nor for β) are referred to as ‘neutral markers’.

**Figure 2:**
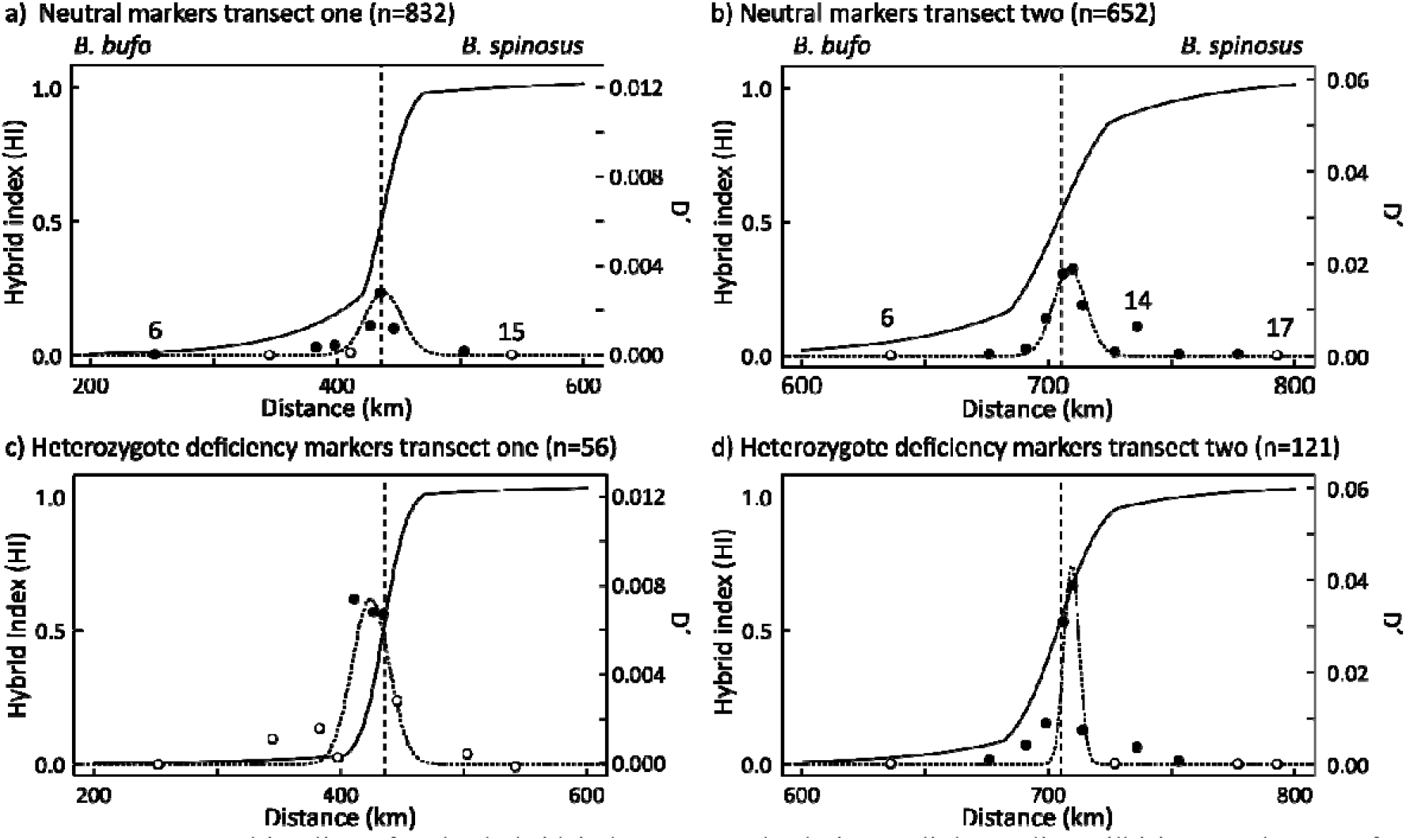
Geographic clines for the hybrid index (HI) and admixture linkage disequilbirium peaks (D’) for neutral markers (i.e. not assigned by BGC as an outlier in any category) according to the genomic clines analysis for (a) transect one and (b) transect two, and for heterozygote deficiency markers (β > 0) for (c) transect one and (d) for transect two. The x-axis shows distance along the transect, note that the axis for transect two is half the length (200 km) of the axis for transect one (400 km). The y-axis on the left shows the HI (solid line), and the y-axis on the right shows D’ (dotted line and dots). The right y-axis for transect one is five times shorter than for transect two. Solid dots show D’ significantly different from zero, whilst open dots are not significantly different from zero, based on 95% confidence intervals.

The HI cline based on neutral markers in transect one (northwest France) shows a high level of asymmetric introgression from *B. spinosus* into *B. bufo* by a fitted cline shape with a tail (Q_left_ = 15.0, Q_right_ = 8.8), whereas transect two (southeast France) shows a pattern of symmetric introgression by a cline shape without tails (Q_left_ = 11.0, Q_right_ = 11.0; Fig. 2, Table S.4, Fig. S.8, Fig. S.9). The cline widths for neutral markers in both transects are similar (47 km and 49 km), and the cline shape in the centre is generally steep (Fig. 3). The HI clines for barrier markers are symmetrical in both transects (Table S.4).

**Figure 3:**
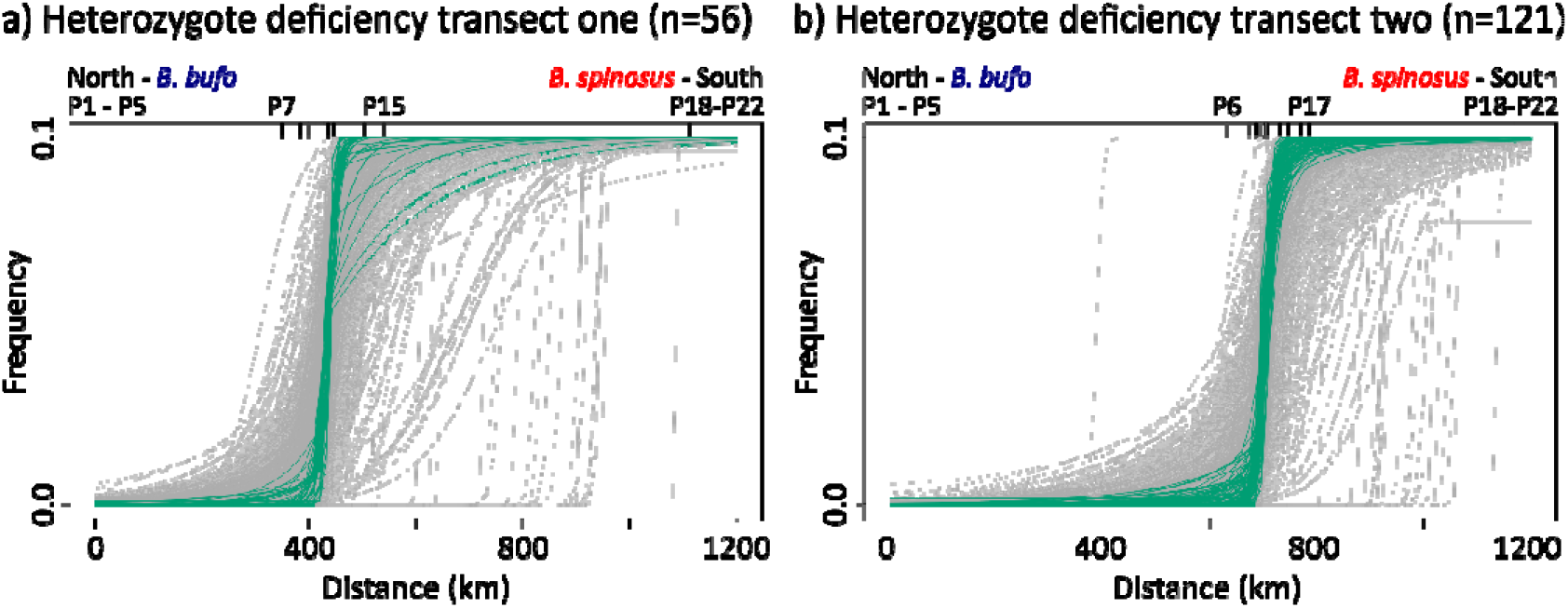
Geographic clines for markers showing barrier markers (green) for (a) transect one and (b) transect two with frequency of the *B. spinosus* allele on the y-axis and distance along the transect on the x-axis. Inward ticks on the top of the graph and notation near inward ticks on the top of the graph (P) refers to locations in Fig. 1.

### Admixture linkage disequilibrium

The admixture linkage disequilibrium (D’) for neutral markers shows a peak in the centre of the hybrid zone in both transects, although the amplitude of the peak in transect one is one fifth of the peak in transect two (Fig. 2). As barrier markers are less likely to flow away from the hybrid zone centre, the peak of D’ for barrier markers may be more sensitive to shifts of the hybrid zone centre. In transect one, barrier markers showed a peak with higher D’ on the *B. bufo* side of the hybrid zone, but displacement of the Gaussian curve was not significant;as the AIC was nearly equal when constraining the peak of D’ to the cline centre (435 km) as when the peak was fitted unconstrained (resulting in a peak at 423 km). In transect two, barrier markers showed a peak of D’ in the centre of the hybrid zone.

The lower peak of D’ in transect one corresponds to a lower number of steep clines observed and underlies the lower estimates of effective selection against hybrids (s*) and lifetime dispersal compared to transect two, and not to a difference in cline width as these are comparable (see results section “geographic cline analysis” above). For transect one s* is 0.0022 (95% CI 0.0012-0.0034) whereas it is about an order of magnitude greater for transect two, where s* is 0.0195 (95% CI 0.0143-0.0252; Table S.5). The s* based on barrier markers for transect one is 0.0101 (95% CI 0.0054-0.0152) and for transect two 0.0344 (95% CI 0.0220-0.0470). Notably, the confidence intervals for both estimates of s* in each hybrid zone do not overlap. We estimated the lifetime dispersal distance based on neutral markers for transect one and two at 1.8 (95% CI 1.3-2.2) and 4.0 (95% CI 3.4-4.5) km per generation, respectively.

## Discussion

### Asymmetry of introgression in northwest France

In northwest France (transect one), Bayesian genomic cline analysis indicates a significant asymmetry in gene flow, with more alleles from *B. bufo* flowing into *B. spinosus*, than the other way around (Table 1). On the other hand, a shift in the HI geographic cline based on neutral markers, which is the baseline to determine outliers in the Bayesian genomic clines analysis, shows asymmetric neutral introgression from *B. spinosus* into *B. bufo* (Fig. 2). The combination of a high amount of possibly selective gene flow from *B. bufo* into *B. spinosus*, and a tail of neutral introgression towards the north (from *B. spinosus* into *B. bufo*) could be pointing to southward hybrid zone movement due to an advantage of *B. bufo* over *B. spinosus*. A neutral tail of introgression towards the north was previously interpreted as a tail of introgression in the wake of (past) hybrid zone movement, and the hybrid zone was thought to have stabilised at a gradient of elevation (Arntzen et al., 2016). The pattern of introgression currently observed, a tail towards the north with a coincident peak of admixture linkage disequilibrium (D’) is highly similar to the pattern observed in van Riemsdijk et al. (2019). Alternative explanations for such asymmetric introgression could be asymmetric reproductive isolation due to mating preferences or mitonuclear incompatibilities.

When studying single transects, often more than one potential explanation for asymmetric introgression across a hybrid zone are reported. Providing solid proof of hybrid zone movement or asymmetric reproductive isolation proves to be difficult (Buggs, 2007; Toews & Brelsford, 2012; Brandvain, Pauly, May, & Turelli, 2014; Wielstra, 2019). Yet some studies provide convincing support for one explanation over the other by combining multiple lines of evidence. For example, in a group of Neotropical jacanas (*Jacana*), females of the larger and more aggressive species more often mother hybrid offspring in sympatric regions, which resulted in a shift of female body mass relative to the genetic cline centre (Lipshutz et al., 2019). In crested newts (*Triturus*) the position of enclaves (distribution relicts), predictions of distribution models, and genome-wide asymmetric introgression support hybrid zone movement as a cause of asymmetric introgression (Wielstra & Arntzen, 2012; Wielstra, Burke, Butlin, & Arntzen, 2017). For the *Bufo* hybrid zone, additional evidence in the form of behavioural, breeding, distribution modelling or simulation studies may thus provide stronger support for the cause of the asymmetric introgression observed in northwest France.

### Symmetry of introgression in southeast France

In southeast France (transect two), both genetic and genomic cline analyses reveal equal amounts of introgression on both sides of the hybrid zone. Therefore, the zone appears to be stable. Whilst the geography of the northwest transect is relatively homogeneous (e.g. altitude, Fig. 1), the hybrid zone in southeast France locally runs in parallel to landscape features (rivers and mountains) that are associated with the species range border (Arntzen et al., 2018). Barrier genes may be coupled to a steep gradient of local adaptive genes between two populations and stabilise a hybrid zone at an ecotone (Bierne, Welch, Loire, Bonhomme, & David, 2011). In such a situation, the local steep environmental gradient may be repeated in the vicinity of the hybrid zone centre (e.g. the presence of multiple hills in a row), but the barrier effect may keep the locally adaptive genes locked in the hybrid zone (e.g. on one hillside, Bierne et al., 2011). The hybrid zone in southeast France may be trapped at such a steep gradient of locally adaptive markers.

### *A barrier to gene flow in* Bufo *transects*

The number of barrier markers (as identified in the genomic cline analysis) in transect one (northwest France; n=56) is approximately half that of transect two (southeast France; n=123), and the estimated selection against hybrids in the neutral markers in the dataset is significantly lower in transect one (s* is 0.0022) than in transect two (s* is 0.0195, Table 1, Table S.5). As expected, the majority of the barrier markers as identified by the genomic cline analysis also show a narrow geographical cline with a transition confined to the centre of the hybrid zone. Our results thus support the idea that the higher the number of barrier genes restricting gene flow is, the higher the overall effective selection against hybrids will be (Barton, 1983; Barton & Gale, 1993; Bierne et al., 2011; Vines et al., 2016). It thus appears that the barrier effect is less prominent in transect one than in transect two.

A difference in negative selection in different parts of the hybrid zones can be caused by several biological processes, including differences in the length of secondary contact, a difference in linkage of barrier genes and involvement of different genetic groups, as we further discuss below. The barrier markers, with relatively steep clines, for which the geographical cline centre is not fixed to the hybrid zone are mostly “shifted” towards the south, from *B. bufo* into *B. spinosus* (Fig. 3). The transition of these few shifted barrier markers is somewhere between the last sample location of the transect and the reference populations, and more samples towards the south would be needed to determine their shape and centre more precisely.

During postglacial expansion, secondary contact between *B. bufo* and *B. spinosus* is suggested to first have established in the southeast of France and at a later point in the northwest (Arntzen et al., 2017), closing the gap between the two species in a zipper-like manner. As a consequence, cline coupling may have progressed further towards reproductive isolation after secondary contact, and is still ongoing throughout the hybrid zone (Harrison & Larson, 2016; Butlin & Smadja, 2018; Dagilis, Kirkpatrick, & Bolnick, 2019). Species divergence with gene flow is predicted to result in more clustering of neutral and barrier loci in certain genomic regions, than species divergence without gene flow, and in this way may provide a stronger barrier against gene flow (Noor, Grams, Bertucci, & Reiland, 2001; Rieseberg, 2001; Emelianov, Marec, & Mallet, 2004; Nosil, Funk, & Ortiz-Barrientos, 2009; Yeaman & Whitlock, 2011; Harrison & Larson, 2016; Rafajlović, Emanuelsson, Johannesson, Butlin, & Mehlig, 2016; Schumer et al., 2018). The barrier markers shared across both transects may therefore be clustered in a few low-recombination regions, or linked to the causal variant and maintained through linkage to loci under strong selection. Whether the higher number of barrier markers observed in the southeast of France is due to the age of the hybrid zone, and more specifically, due to the presence of a higher number of barrier genes, or lower recombination rate surrounding barrier genes is not possible to determine with the current dataset, without the aid of a linkage map or sequenced genome.

The two transects share 26 barrier markers (the union of 56 and 123 barrier markers identified in each transect), which is unlikely to be the result of chance (see statistic test in Materials & Methods, and details in Table 1). Overlap of markers involved in the restriction of gene flow in multiple transects pointed to intrinsic similarities of a barrier effect in the sunflower (*Helianthus*) hybrid zone, one of the first hybrid zones studied with multiple transects, with gene flow restricted by genomic regions linked to pollen sterility and chromosomal rearrangements (Rieseberg, Whitton, & Gardner, 1999; Buerkle & Rieseberg, 2001). Under laboratory conditions the barrier genes were also found to restrict gene flow, which excluded the possibility of the involvement of external factors (Buerkle & Rieseberg, 2001). In field crickets (*Gryllus*) the same barrier genes restricted gene flow in two sections of the hybrid zone and barrier genes were linked to intrinsic factors of prezygotic isolation (Larson, Andrés, et al., 2013; Larson, Guilherme Becker, Bondra, & Harrison, 2013; Larson et al., 2014). In both these examples, the overlap of barrier genes could be linked to processes important for reproduction, which appear to have played a role in the initial isolation of two species.

However, more often than not, the markers restricting introgression differ between transects in the same hybrid zone (see for an overiew Harrison & Larson, 2016). In the well-studied house mouse hybrid zone (*Mus*), patterns of restricted gene flow differ among transects, suggesting also different genetic architectures of isolation between the two species (Teeter et al., 2009). However, just as for *Bufo*, there were markers which show restricted introgression in both transects in *Mus*, which were linked to hybrid sterility in laboratory settings (Janoušek et al., 2012). A similar situation is observed in the *Bufo* hybrid zone. Experimental studies are required to explore the functional roles of the barrier markers in the *Bufo* hybrid zone.

### *Intraspecific divergence within* B. bufo

The *B. bufo* toads in northwest France possess an mtDNA clade that resided in a refugium in the northern Balkans during the last glacial maximum (22,000 BP), whilst *B. bufo* in southeast France carry mtDNA variants that survived in Italy, and in the northern and western Balkans (Garcia-Porta et al., 2012; Recuero et al., 2012; Arntzen et al., 2017). Meanwhile, *B. spinosus* individuals from France belong to a lineage derived from a single Iberian refugium (Garcia-Porta et al., 2012; Recuero et al., 2012; Arntzen et al., 2017). The difference in distance from these refugia to the current position of the hybrid zone may also have resulted in a later point of contact in northwest France, compared to southeast France and, perhaps, reduced opportunity for secondary contact in the northwest during previous interglacials. Additionally, *B. bufo* shows structural differences in chromosome morphology across its range, with all homogametic chromosomes observed in *B. bufo* in Russia (which presumably belong to the same mitochondrial clade occurring in northwest France), and a heterogametic pair of chromosomes in female *B. bufo* in Italy, whereas chromosome morphology within *B. spinosus* appears to be uniform (Morescalchi, 1964; Birstein & Mazin, 1982; Pisanets et al., 2009; Skorinov et al., 2018). Intraspecific mitochondrial and chromosome divergence in *B. bufo* may thus be reflected by (other) nuclear genetic substructure. Yet, the Structure and PCA results based on the total dataset, despite the inclusion of all 4,863 RAD markers, did not indicate the presence of multiple genetic groups in our *B. bufo* samples. But, given our sampling scheme (Fig. 1), this could be due to the lack of (pure) parental populations belonging to such potential genetic groups (Rogers & Bohlender, 2015). A range-wide phylogeographic study based on nuclear DNA is required to assess whether diverged (nuclear) *B. bufo* groups are involved in the hybrid zone.

### Evolution of hybrid zones

Long hybrid zones, such as the common toad and house mouse hybrid zones, appear to have two evolutionary trends in common; 1) barrier markers shared between transects, reflecting an overall barrier to gene flow along the entire hybrid zone, and 2) barrier markers specific to individual transects. Hence, long hybrid zones can provide unique insights in the different ways two lineages move towards speciation, or in the opposite direction, towards complete merging (Teeter et al., 2009; Larson, Andrés, et al., 2013; Harrison & Larson, 2014; Larson et al., 2014). The current study shows that the process of cline coupling, where additional barrier genes are recruited and converge geographically towards the hybrid zone (Butlin & Smadja 2018), may result in spatial variation in the set of barrier markers employed along the length of the hybrid zone.

## Conclusion

We find an overlap of barrier markers between two widely separated transects in the *Bufo* hybrid zone, which indicates that a range wide barrier effect has evolved. The barrier effect is strong enough to have prevented the two *Bufo* species from merging despite secondary contact having been established about 8000 years ago (Arntzen et al., 2016; Arntzen, 2019). However, we propose that potential genetic substructure within *B. bufo* complicates the interpretation of overlap and differences between transects within this hybrid zone, and we recommend that future research explores the presence of subgroups based on genome-wide nuclear DNA data for a wider geographic range. The generation of a high-density linkage map or reference genome will be helpful to infer patterns of linkage and barrier loci in more detail. Laboratory crosses of individuals from the resulting intraspecific *B. bufo* groups and *B. spinosus* could verify potential modes of (asymmetric) reproductive isolation (e.g. Malone & Fontenot, 2008; Stöck et al., 2013; Brandvain et al., 2014). The *Bufo* hybrid zone provides an excellent opportunity to separate a general barrier to gene flow from local reduction in gene flow specific to individual transects.

## Acknowledgements

We thank Frido Welker, Tara Luckau, Roger K. Butlin, the members of the Butlin laboratory at the University of Sheffield, and the Allentoft laboratory at the Natural History Museum in Copenhagen for support and discussion. The PhD position of IvR is supported by the ‘Nederlandse Organisatie voor Wetenschappelijk Onderzoek’ (NWO Open Programme 824.14.014). This project has received funding from the European Union’s Horizon 2020 research and innovation programme under the Marie Skłodowska-Curie grant agreement No. 655487. Part of this project was carried out by IR at the Shaffer laboratory in Los Angeles, at the University of California. This study trip has been sponsored by the Leiden University Fund / Swaantje Mondt Fonds (D7102). MR was funded by the Hasselblad Foundation Grant to Female Scientists, a grant from the Swedish Research Council Formas, and by additional grants from Swedish Research Councils (Formas and VR) to the Centre for Marine Evolutionary Biology at the University of Gothenburg (www.cemeb.science.gu.se)

## Data Accessibility Statement

In- and output of analyses, and custom R scripts will be available at Dryad. Raw sequencing data will be available at GenBank.

## Author contributions

IvR, JWA, BS, and BW designed the study. IvR and JWA collected samples. IvR performed the laboratory work and data assembly with contributions from GB, EMM, PS, and ET. IvR analysed and interpreted the data with contributions from BS, BW, JWA, MR, and PS. IvR wrote the manuscript with input from all authors.

