## Supplementary figures for "Spatial variation in introgression along a toad hybrid zone in France"

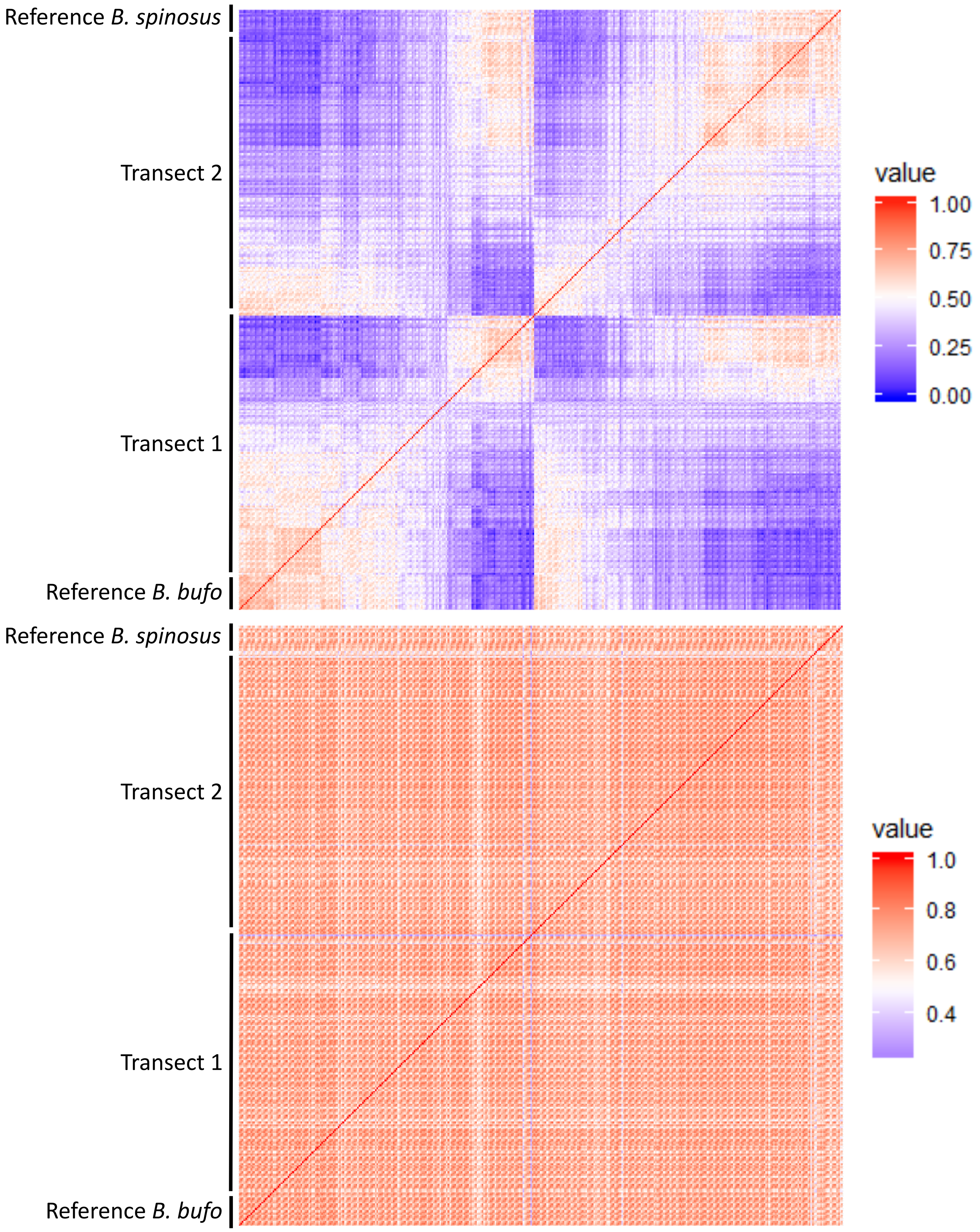


**Figure S.1**: Missingness matrices comparing the amount of number of loci missing between two individuals. The top graph shows the 50% missing data with on the right the colour range for missing values from 0-1, and on the bottom the 90% missing data with a colour range from 0.2-1. A value of 1 indicates no missing data, and 0 all data is missing.

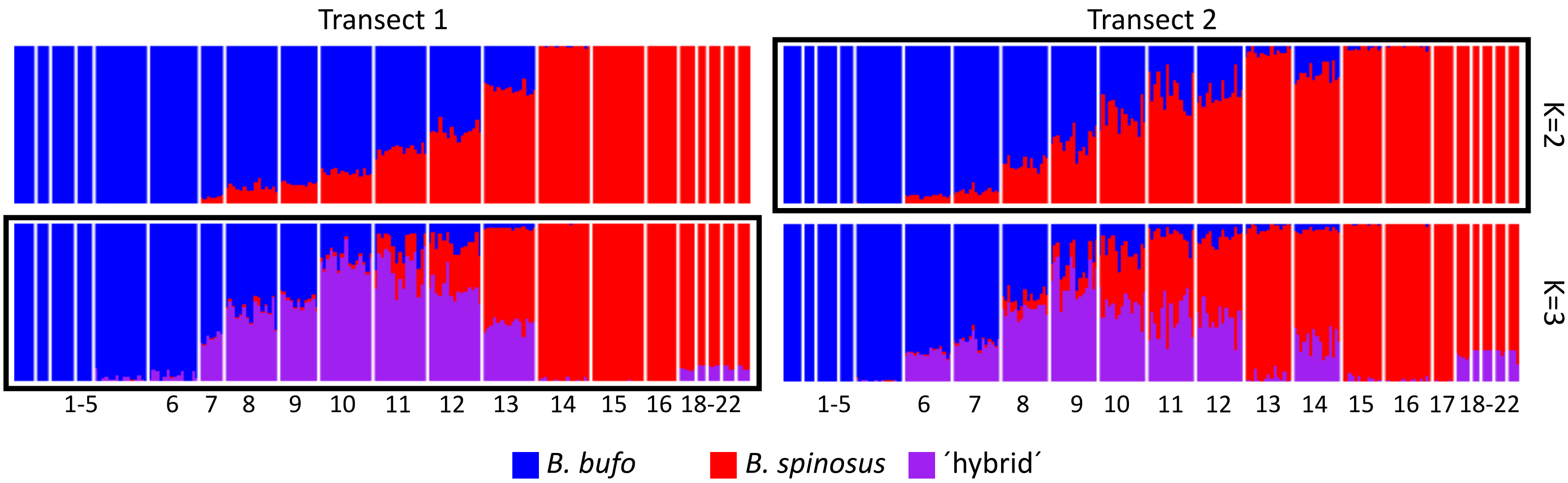


**Figure S.2**: Structure plots for transect 1 and 2 based on the 50p dataset for two genetic clusters (K=2) and three genetic clusters (K=3), with best K for each transect indicated with a black square around the plot. Blue bars indicate the genetic cluster identified as *Bufo bufo*, red bars indicate the genetic cluster identified as *B. spinosus* and purple bars indicate the genetic cluster identified as ´hybrids´.

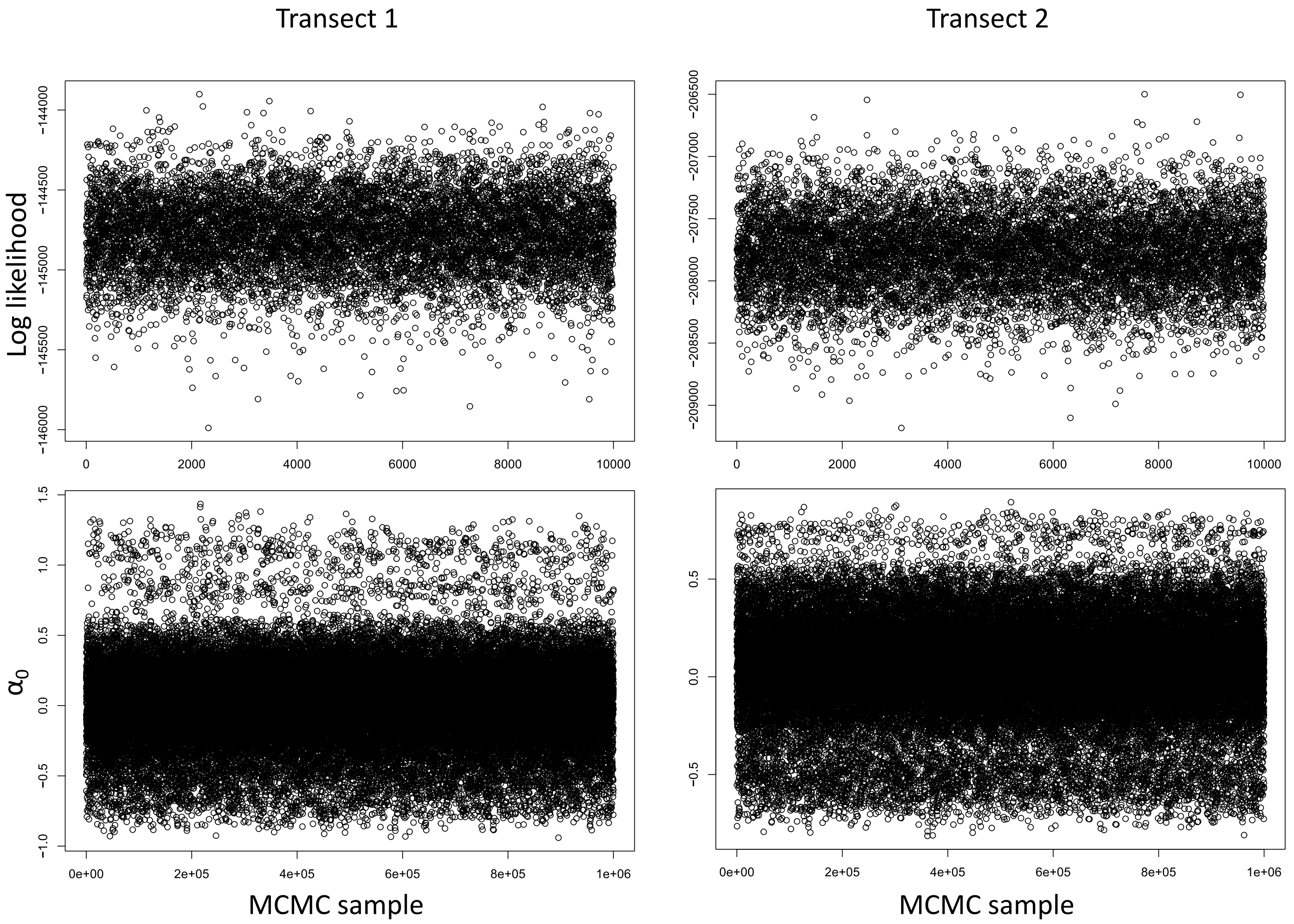


**Figure S.3**: Scatter plots showing the log-likelihood and $\alpha_0$ as a fuction of MCMC sample. On the left are the plots for transect 1, and on the right for transect 2. The bottom plots represent a random selection of 10% of the total samples generated. For both transects, a single MCMC chain was run for 75,000 steps and sampleswere taken from the posterior distrubution every $5^{\text{th}}$ step following a burn-in of 25,000 steps.

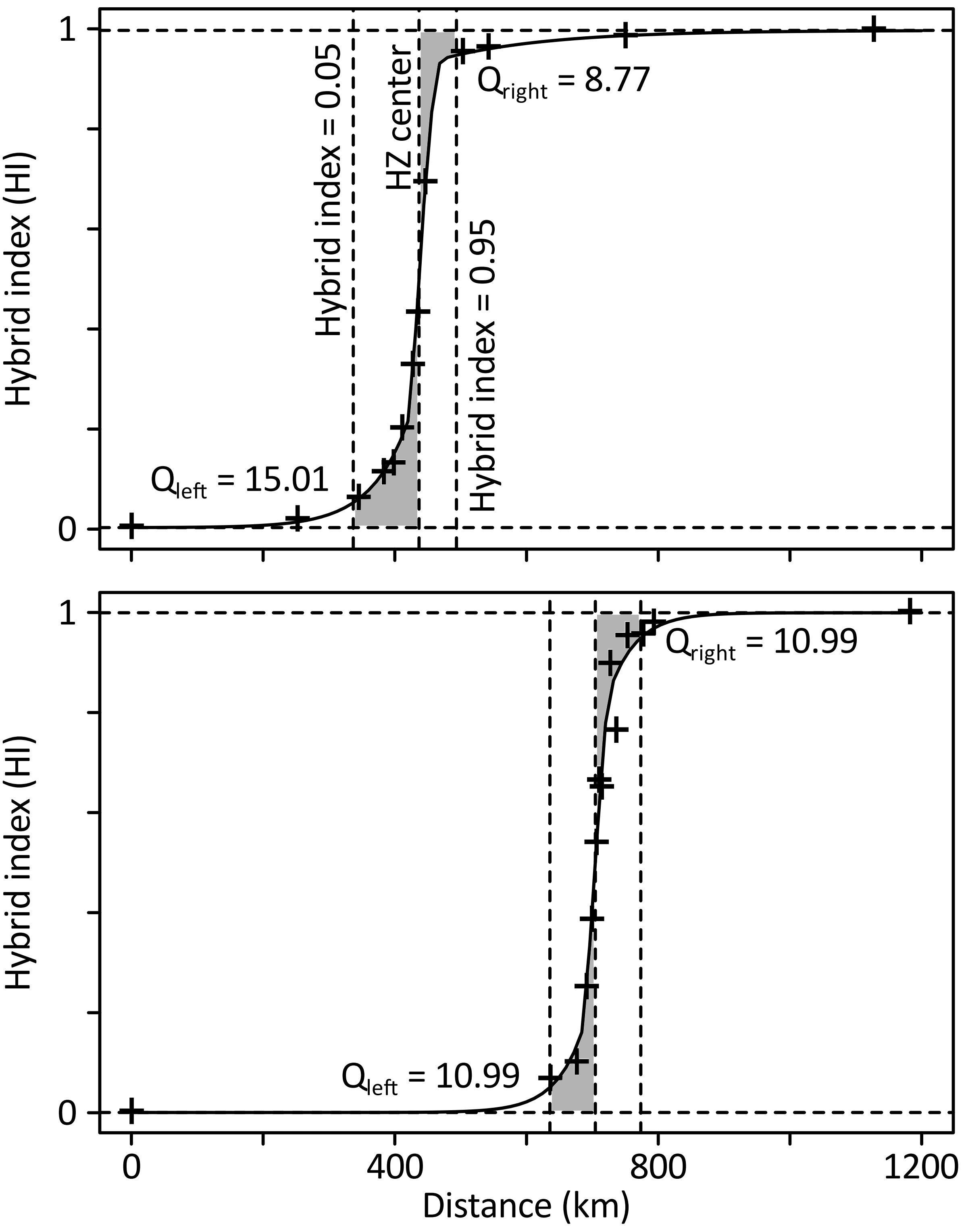


**Figure S.4**: Visualisation of caclucation of the area under the curve for geographic clines. This example figure is based on the hybrid index (HI) clines for neutral markers for transect 1 (top), and transect two (bottom), therefore, instead of gene frequency, as would be the case in cline fittings for single markers, the y axis shows the HI. The x axis shows the distance in kilometers (km). The ´+´ sign shows the actual datapoints the cline was fitted to. For each cline, the cline centre is taken to represent the hybrid zone (HZ) centre, and the cline formula is used to calculate at which distance the hybrid index (or frequency) would be 0.05 (left border) or 0.95 (right border). Using the integral() function from the R package pracma, the area under the curve between the left border and the HZ center is calculated ($Q_{left}$). The area above the curve between the HZ center and the right border is calculated by substracting the area under the curve from the total area on the right side:

$Q_{right} = 1 * ( \text{right border} - \text{HZ center} ) - Q_{right\_under}$

The calculations were made with a custom R script.

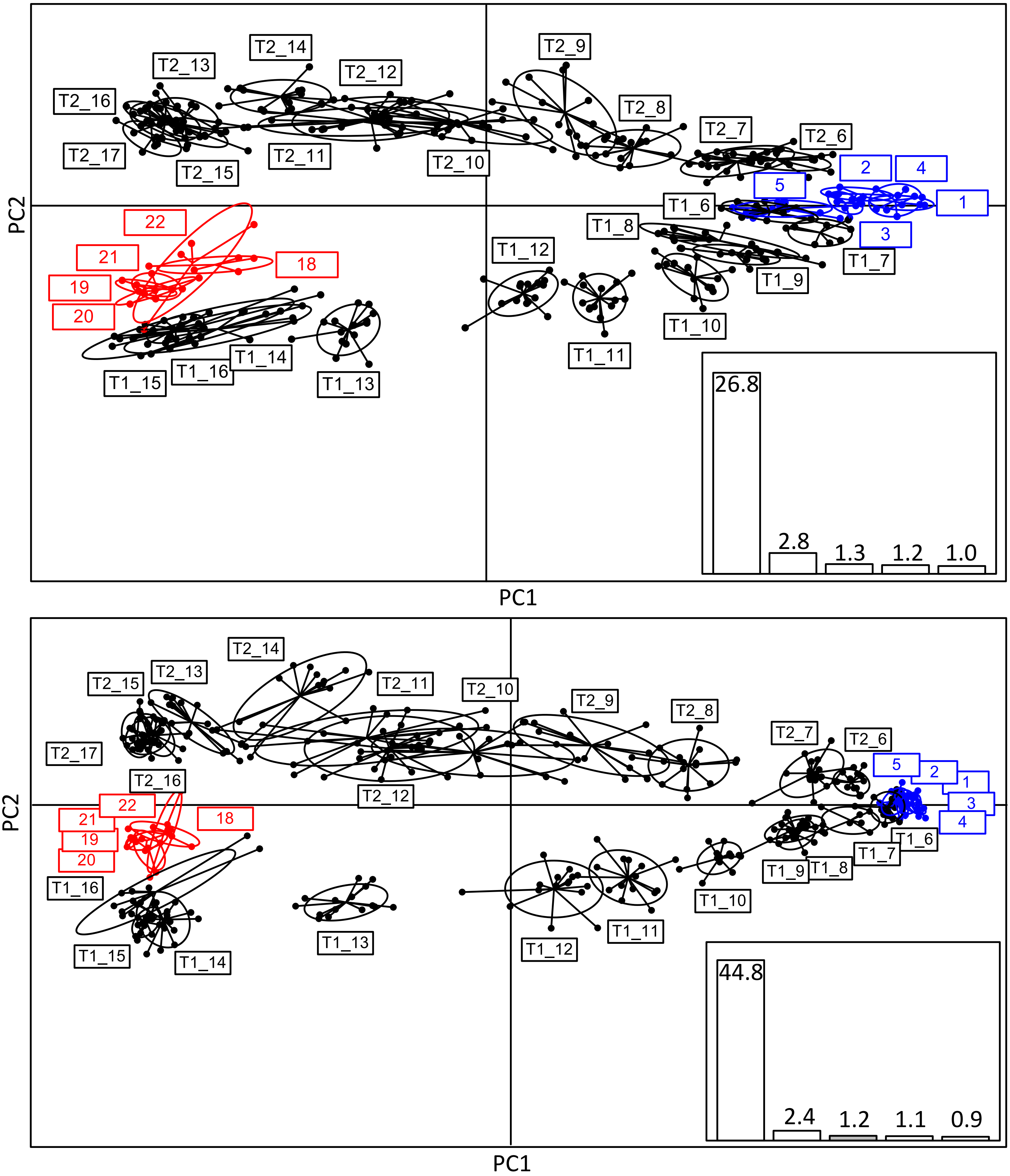


**Figure S.5**: Principal component analysis of the full 50% dataset (top) and full 90% dataset (bottom) taking a random SNP from each of the 4,863 and 986 markers, respectively. The first axis (PC1) for both plots appears to represent geography from south (left, *Bufo spinosus*) to north (right, *B. bufo*). The reference populations of *B. bufo* were coloured blue and the reference populations for *B. spinosus* red. Populations which were regarded as transect populations are black (T1 = transect 1, T2 = transect 2). The inset shows the eigenvalues for the first 5 components.

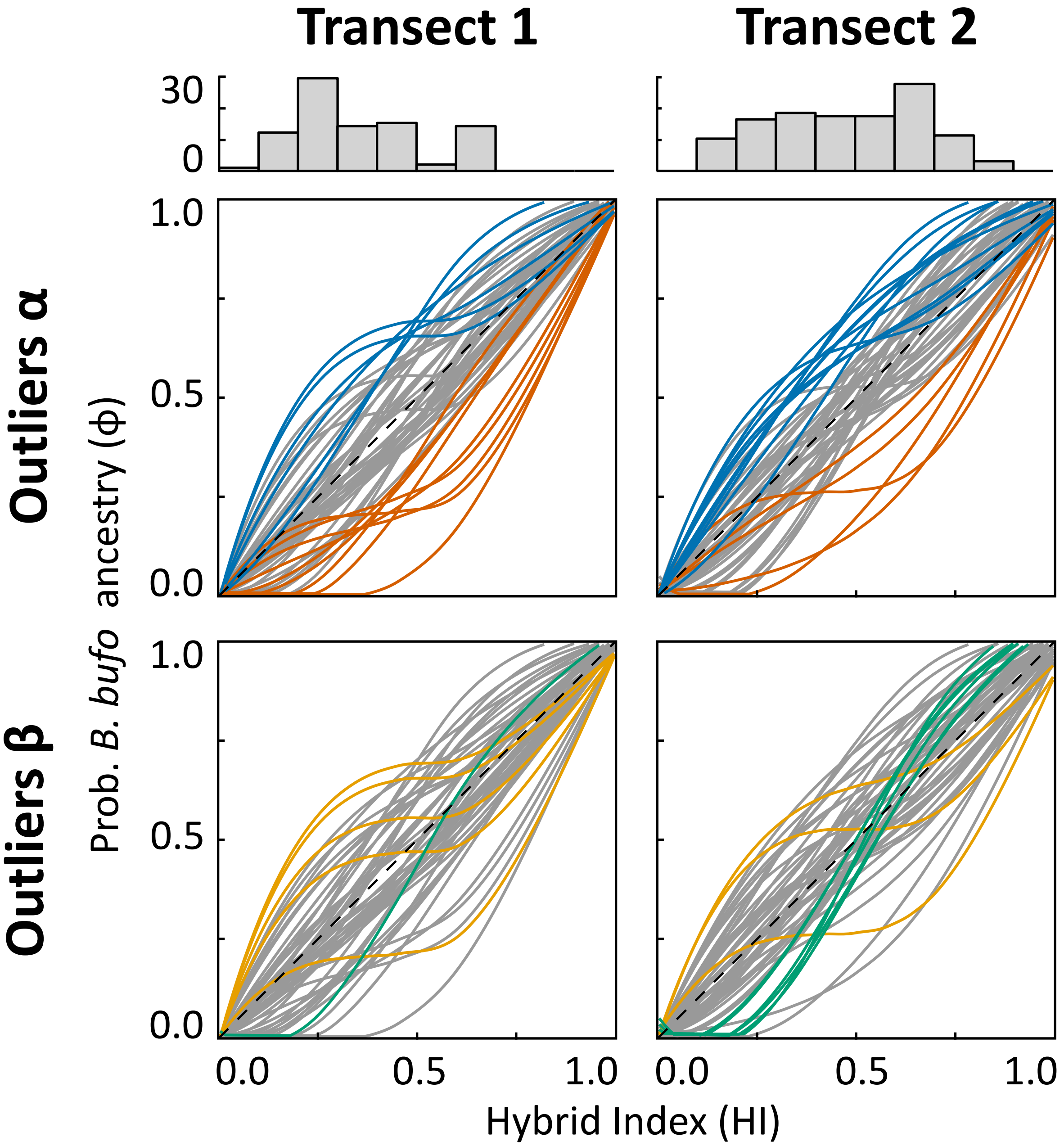


**Figure S.6**: Bayesian genomic clines for transect one left and for transect two right, with on top a bar graph with the number of individuals per bin of hybrid index (0-1 with bin width 0.1). The clines show a sample of the first 50 markers from the total dataset (1,189 markers). A cline is considered to be an outlier when the 99.9 % confidence interval (CI) dos not include 0. The top two panels show outliers for $\alpha > 0$ (blue lines) and $\alpha < 0$ (dark orange lines), and the bottom two panels show outliers for $\beta > 0$ (green lines) and $\beta < 0$ (orange lines).

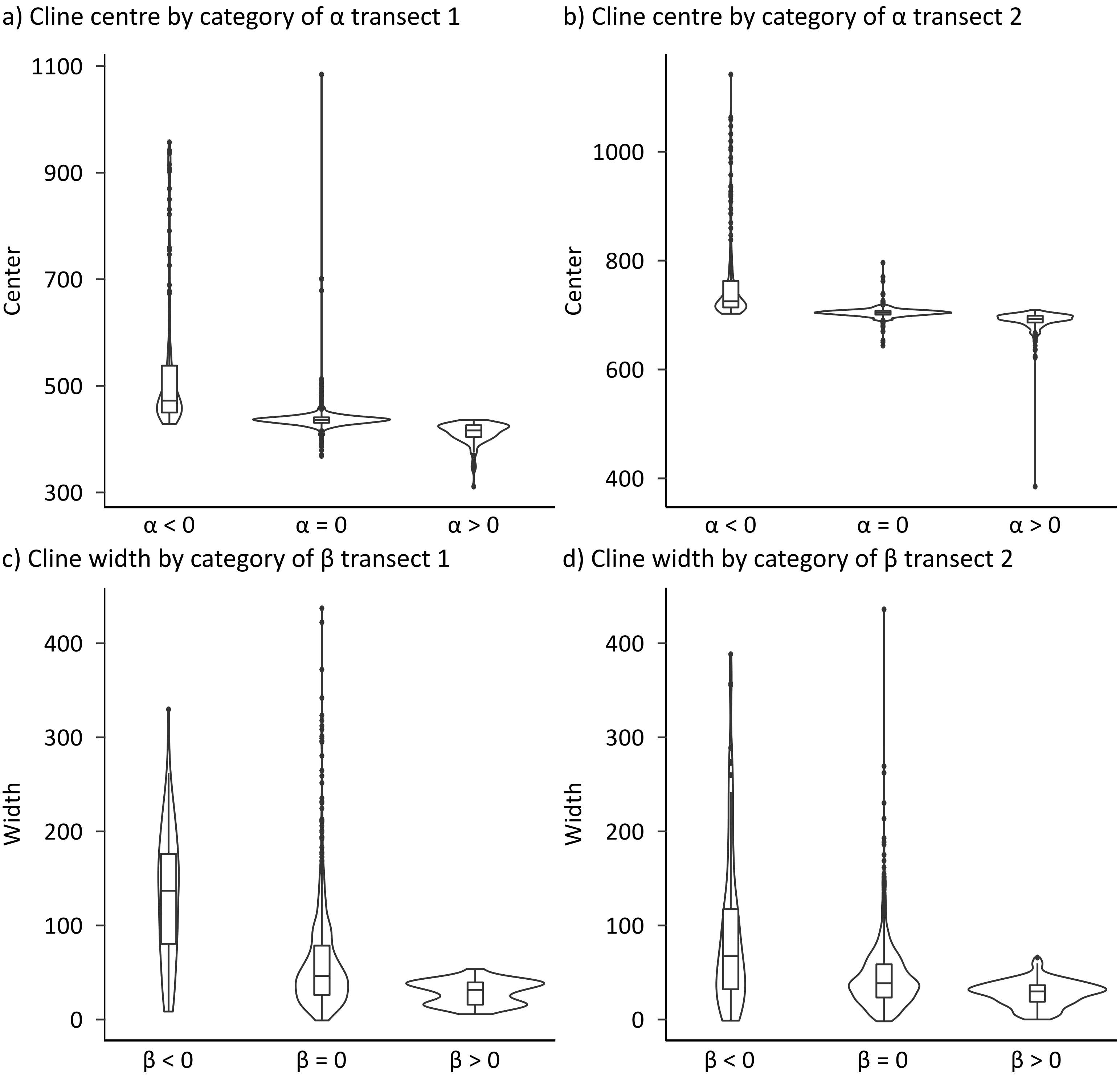


**Figure S.7**: Correlation between outlier categories from the Bayesian genomic clines (BGC) analysis and the cline parameters center and width obtained from geographic cline analysis (HZAR).

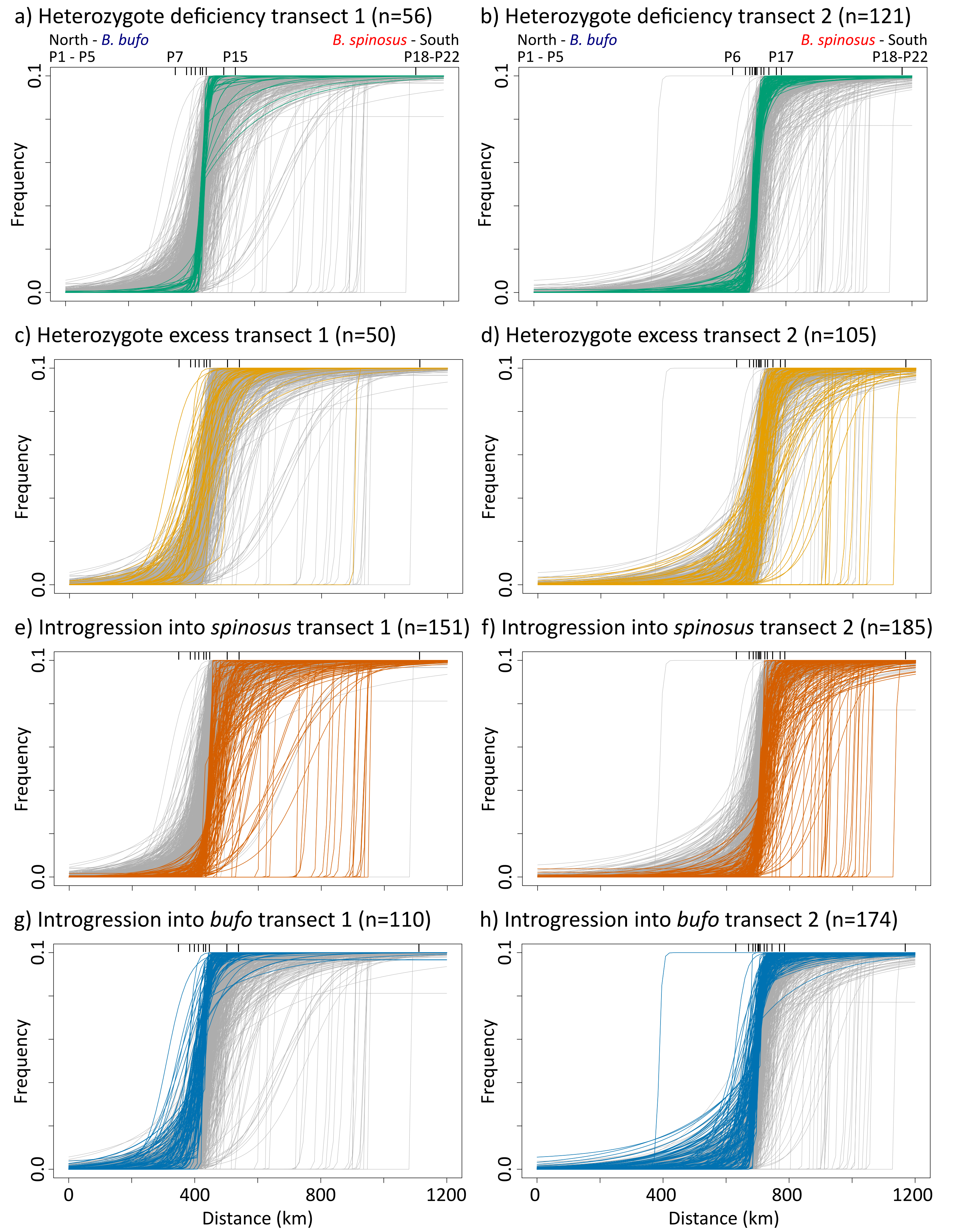


**Figure S.8**: Geographic clines for all markers (grey) and different Bayesian genomic cline (BGC) outliers for transect 1 (a, c, e, g) and transect 2 (b, d, f, h), with frequency of the *Bufo spinosus* allele on the y-axis and distance along the transect on the x-axis. Heterozygote deficiency markers (β > 0, green) are geographically restricted, whilst heterozygote excess markers are unrestricted (β < 0, light orange). Clines with directional introgression from *B. bufo* into *B. spinosus* are situated left of the hybrid zone center (α < 0, dark orange), whilst clines with directional introgression from *B. spinosus* into *B. bufo* are situated left of the hybrid zone (α > 0, blue). Populations are indicated by inward ticks on the top of the graph, and notation refers to population numbers on the map.

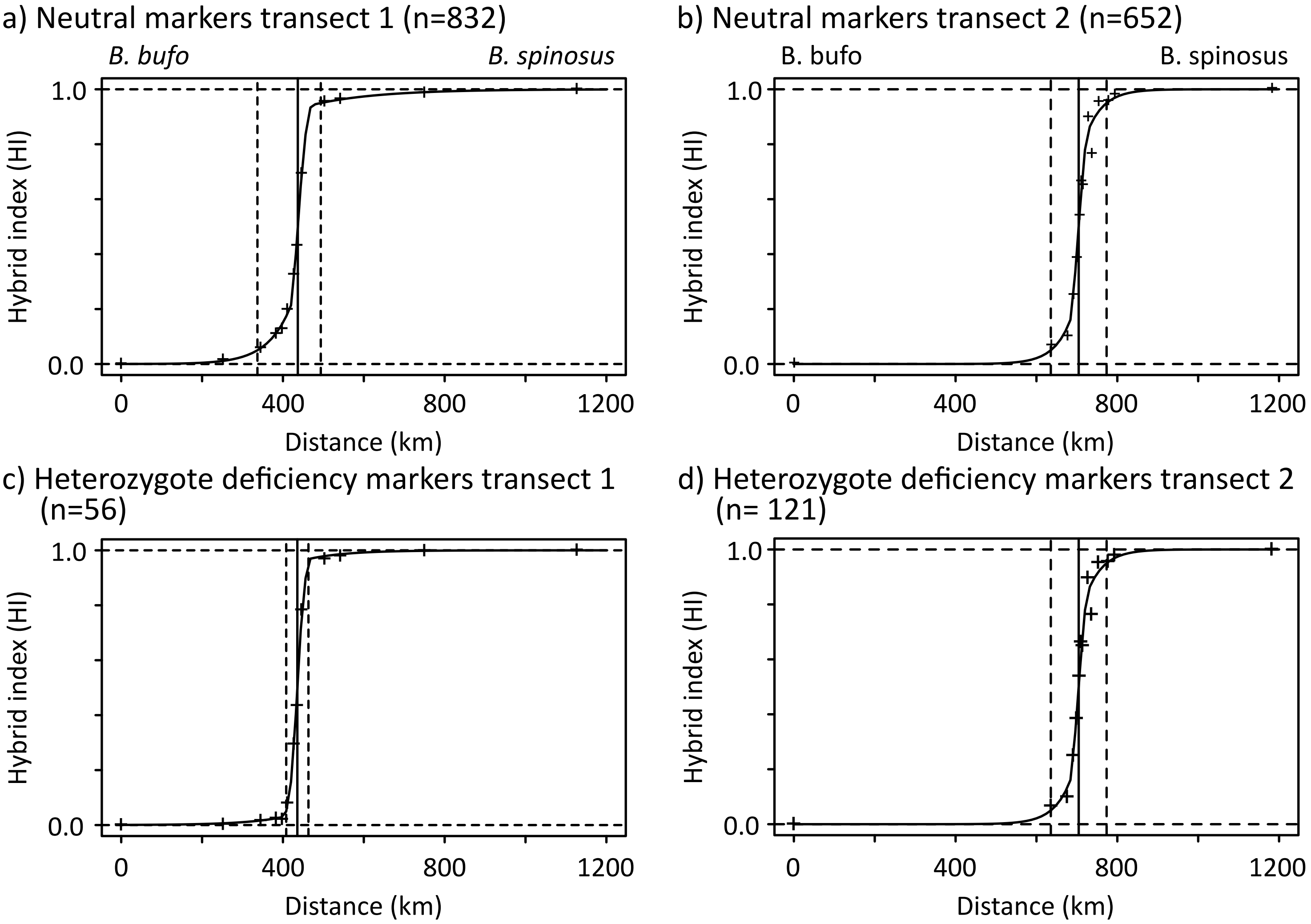


**Figure S.9**: Geographic clines for (a) neutral markers in transect 1, (b) neutral markers in transect 2, (c) heterozygote deficiency markers in transect 1, and (d) heterozygote deficiency markers for transect 2. The x-axis shows distance, the y-axis shows the hybrid index. The vertical solid line shows the cline centre, the vertical and horizontal dashed lines show the cut offs of a hybrid index of 0.05 and 0.95, and the value 0 and 1, which are used to determine the area underneath the curve. Actual hybrid index for each population is indicated with ´+´.
