## Appendix 1 for "Spatial variation in introgression along a toad hybrid zone in France"

**Appendix 1: Library duplicates**

Libraries for six individuals (three from a *B. bufo* reference population, three from a *B. spinosus* reference population) were prepared in triplicate, see Table 1. The first library (“_A”) was prepared in a separate plate from the last two libraries (“_B” and “_C”). All triplicate libraries were prepared with separate iTru primers to differentiate the library preparations after assembly. Libraries were treated as separate samples until the end of the assembly with ipyrad (Eaton, 2014). With the subsequent data, we calculated the overall error rate introduced during library preparation, PCR, sequencing, assembly, and SNP calling.

To calculate the error rate we used a custom R script, included in the online supplements. We used the full SNP file generated by ipyrad from the 50% dataset in phylip format. We converted this to a FASTA before reading the data into R. Each sequence was compared with the other sequences within each species. The results are shown in Figure 1. On top are the heatmap tables with the percentage difference caused by missing data (represented by either “-“ or “N” in the FASTA sequence), on the bottom are the heatmap tables with the percentage of difference in actual nucleotide call (represented by e.g. both a “T” and an “A” in the same position). We found that most of these calls include errors in calling of a heterozygote (represented by IUPAC codes) and calling a homozygote of one of those nucleotides, which are mostly dependent on read depth. In general we thus underestimate the number of heterozygotes. The mean error rate is therefore estimated to be 0.49 %. The library with the lowest total amount of missing data (represented by “-“ or “N”) were used in the downstream analyses to represent the individual (indicated with “ ^+^ ”).

**Table 1.** Identification of individuals used in the triplicate experiment, with identification number and “_A”, “_B”, “_C” to indicate the first, second, and third repeat of the library preparation. The individuals with the lowest total amount of missing data (represented by “-“ or “N”) was used in the downstream analyses to represent the individual (indicated with “ ^+^ ”).

| **Individual** | **Species** | **Missing data** |
| --- | --- | --- |
| 1572_A^+^ | *B. bufo* | 17,481 |
| 1572_B | *B. bufo* | 20,576 |
| 1572_C | *B. bufo* | 20,191 |
| 1573_A^+^ | *B. bufo* | 16,344 |
| 1573_B | *B. bufo* | 21,435 |
| 1573_C | *B. bufo* | 20,987 |
| 1574_A^+^ | *B. bufo* | 17,640 |
| 1574_B | *B. bufo* | 23,796 |
| 1574_C | *B. bufo* | 19,183 |
| 4112_A^+^ | *B. spinosus* | 14,937 |
| 4112_B | *B. spinosus* | 22,568 |
| 4112_C | *B. spinosus* | 16,588 |
| 4113_A | *B. spinosus* | 15,860 |
| 4113_B^+^ | *B. spinosus* | 15,508 |
| 4113_C | *B. spinosus* | 19,288 |
| 4114_A | *B. spinosus* | 18,809 |
| 4114_B^+^ | *B. spinosus* | 18,049 |
| 4114_C | *B. spinosus* | 20,999 |

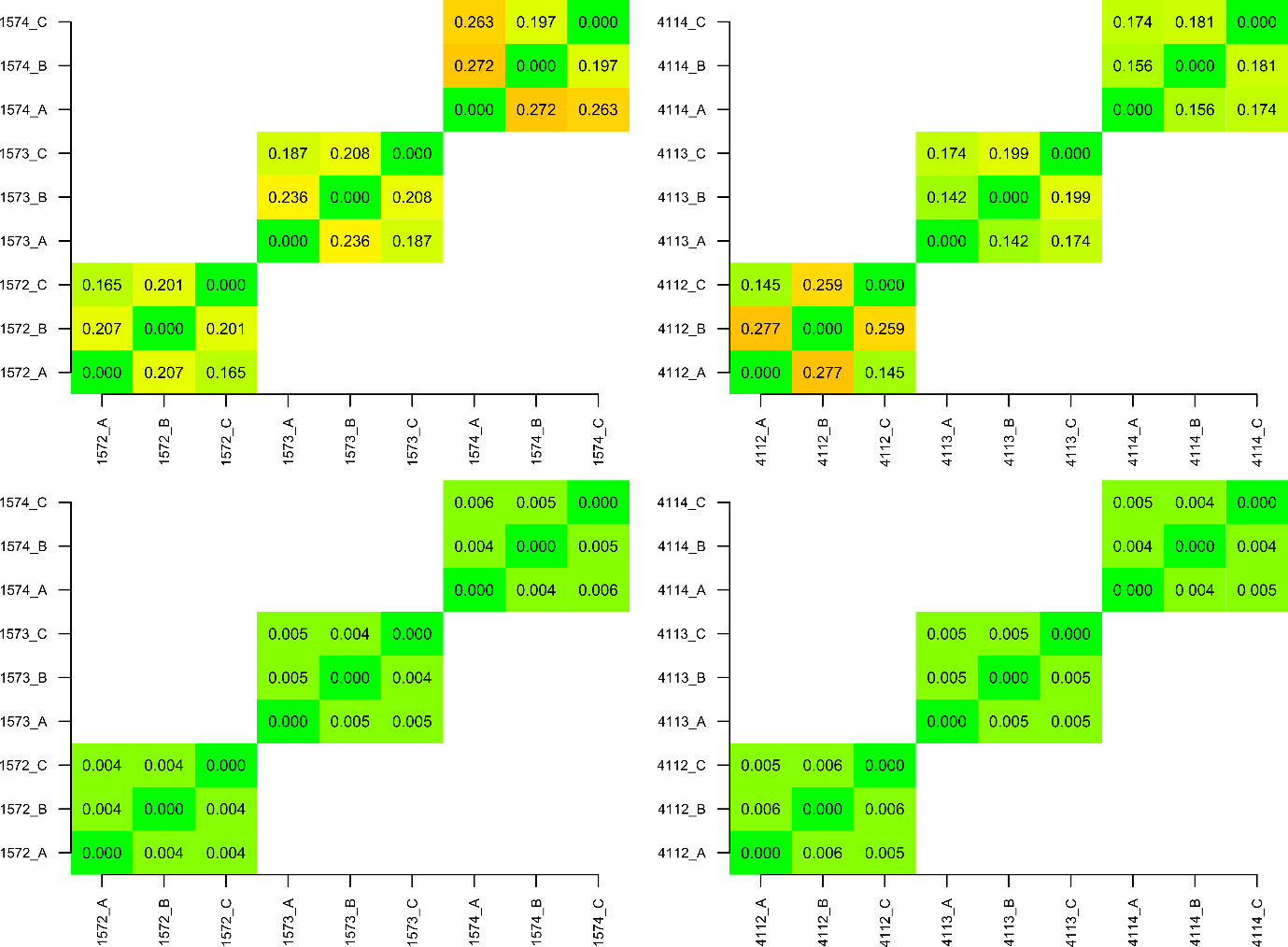

**Figure 1.** Top panels show missing data (represented by either “-“ or “N” in the FASTA sequence), on bottom panels show percentage of difference in actual nucleotide call (represented by e.g. “T” instead of “A”). Left two panels are the *B. bufo* sequences, right panel the *B. spinosus* sequences. Colour scheme is from green (0 missing data) to red (0.9 missing data).
