## Supplemental Data 1 for "Spatial variation in introgression along a toad hybrid zone in France"

**Appendix 2. Bead based clean-up**

This is a protocol to purify ligated stubs from your library, as executed at the Shaffer laboratory.

Prepare:

Take SPRI aliquot and tubes (50 mL) covered in aluminium from fridge 1h before PCR is done, to get to room temperature. Shake tubes with the beads aliquot until entirely dissolved. Prepare ‘boats’ with dH_2_O and freshly prepared EtOH (80%).

Step by step protocol:

1. Add 30 µL of dH_2_O (25 µL when library is prepped) to each library
2. Add 60 µL of beads to each library (e.g. using spreader pipette). Pipet up and down to homogenise
3. Incubate for 4 minutes at room temperature and shortly spin down, this allows your DNA to attach to the beads
4. Capture your beads on a plate magnet for 4 minutes
5. Discard the supernatant (150 µL) whilst the plate remains on the magnet, avoid touching the pellet
6. Add 150 µL of EtOH, wait for 30 seconds, and keep the plate on the magnet, this cleans the DNA
7. Discard the supernatant, take the plate off the magnet, dry beads (about 7 minutes), or until they start to appear cracked or less shiny
8. Add 20 µL of dH_2_0 and pipet up and down to suspend, shortly spin down, and wait for 2 minutes to allow DNA to detach from the beads
9. Move the plate back on the magnet and wait 4 minutes
10. Move, without touching the pellet, the supernatant to a new plate or low retention tubes, these now contain the cleaned library
